# Calcineurin contributes to RNAi-mediated transgene silencing and small interfering RNA production in the human fungal pathogen *Cryptococcus neoformans*

**DOI:** 10.1101/2023.07.25.550548

**Authors:** Vikas Yadav, Riya Mohan, Sheng Sun, Joseph Heitman

## Abstract

Adaptation to external environmental challenges at the cellular level requires rapid responses and involves relay of information to the nucleus to drive key gene expression changes through downstream transcription factors. Here, we describe an alternative route of adaptation through a direct role for cellular signaling components in governing gene expression via RNA interference-mediated small RNA production. Calcium-calcineurin signaling is a highly conserved signaling cascade that plays central roles in stress adaptation and virulence of eukaryotic pathogens, including the human fungal pathogen *Cryptococcus neoformans*. Upon activation in *C. neoformans*, calcineurin localizes to P-bodies, membrane-less organelles that are also the site for RNA processing. Here, we studied the role of calcineurin and its substrates in RNAi-mediated transgene silencing. Our results reveal that calcineurin regulates both the onset and the reversion of transgene silencing. We found that some calcineurin substrates that localize to P-bodies also regulate transgene silencing but in opposing directions. Small RNA sequencing in mutants lacking calcineurin or its targets revealed a role for calcineurin in small RNA production. Interestingly, the impact of calcineurin and its substrates was found to be different in genome-wide analysis, suggesting that calcineurin may regulate small RNA production in *C. neoformans* through additional pathways. Overall, these findings define a mechanism by which signaling machinery induced by external stimuli can directly alter gene expression to accelerate adaptative responses and contribute to genome defense.

**Article summary:** Signaling cascades primarily drive responses to external stimuli through gene expression changes via transcription factors that localize to the nucleus and bind to DNA. Our study identifies an alternative mechanism whereby calcineurin, a key and direct downstream effector of calcium signaling, is involved in post-transcriptional regulation of gene expression through RNAi-mediated small RNA production. We propose that such signaling allows cells to bypass the requirement for communication to the nucleus and rapidly drive stress responses in a reversible fashion.

## Introduction

RNA interference (RNAi) is a mechanism by which cellular gene expression is suppressed post-transcriptionally through targeted RNA cleavage and degradation. RNAi has been proposed to have evolved as a genome defense mechanism against invading RNA viruses and transposons (Buchon and Vaury 2006; Obbard *et al*. 2009). The basic process of RNAi involves either the identification of double-stranded RNA or the generation of double-stranded RNA molecules from single-stranded RNA via RNA-dependent RNA polymerases (Shabalina and Koonin 2008; Obbard *et al*. 2009; Castel and Martienssen 2013). The double-stranded RNA moiety is recognized by Dicer and cleaved into smaller RNA fragments, which are then bound by Argonaute and serve as a template to identify longer target RNA molecules. As a result, the target RNA molecule is destroyed rendering it non-functional and resulting in suppression or silencing of gene expression. RNAi has also been utilized as an important tool to study gene functions by perturbing gene expression levels (Boutros and Ahringer 2008; Castel and Martienssen 2013).

While there are differences in the accessory proteins that operate during RNAi, the core proteins and key steps are highly conserved across evolution from yeasts to humans (Shabalina and Koonin 2008; Lax *et al*. 2020). In fungi, the RNAi machinery has been well characterized in both model and pathogenic species since its initial discovery in *Neurospora crassa* (Cogoni and Macino 1999; Nakayashiki 2005; Nicolas and Garre 2016). In addition to its conventional genome defense role, RNAi contributes to genome integrity by establishing centromere identity, maintaining centromere structure, and driving antifungal drug resistance through epimutations as well as by controlling transposition (Volpe *et al*. 2003; Folco *et al*. 2008; Calo *et al*. 2014; Yadav *et al*. 2018; Chang *et al*. 2019; Priest *et al*. 2022).

*Cryptococcus neoformans* is a basidiomycetous yeast and a human fungal pathogen that primarily causes infections in immune-compromised patients and accounts for ∼20% of HIV/AIDS-related deaths (Rajasingham *et al*. 2017; Zhao *et al*. 2019; Rajasingham *et al*. 2022). RNAi is required for genome defense in *C. neoformans* by silencing both transposons and transgene arrays (Janbon *et al*. 2010; Wang *et al*. 2010; Dumesic *et al*. 2013). Transgene silencing occurs at ∼50% frequency during sexual reproduction and at a lower frequency (0.1%) during mitotic growth and has been termed sex-induced silencing (SIS) or mitotic-induced silencing (MIS), respectively (Wang *et al*. 2010; Wang *et al*. 2012). Studies on the RNAi-mediated silencing mechanism identified several unconventional proteins as part of the RNAi machinery in *C. neoformans* (Feretzaki *et al*. 2016; BURKE *et al*. 2019).

Localization of core RNAi components by direct fluorescence of epitope-tagged proteins revealed their localization at processing bodies (P-bodies), membrane-less organelles involved in RNA processing (Wang *et al*. 2010). Interestingly, previous studies demonstrated that the heat-stress-activated phosphatase calcineurin also localizes to P-bodies during 37°C heat stress in *C. neoformans* (Kozubowski *et al*. 2011). Calcineurin is a heterodimeric phosphatase comprised of the catalytic A and regulatory B subunits (Rusnak and Mertz 2000; Park *et al*. 2019; Ulengin-talkish and Cyert 2023). Calcineurin is activated upon calcium influx in the cell by calcium-bound calmodulin and calcium binding to calcineurin B (Yadav and Heitman 2023). Calcineurin is also the target of two immunosuppressive drugs, FK506 and cyclosporin A, and its activity in *C. neoformans* is inhibited by these drugs. Previous studies have established that calcineurin is essential for both growth at 37°C and sexual reproduction in *C. neoformans* (Odom *et al*. 1997; Cruz *et al*. 2001; Fox *et al*. 2001). Phosphoproteome studies identified several candidate calcineurin substrates localized in P-bodies and explored their roles in thermotolerance and sexual reproduction (Park *et al*. 2016). Specifically, these substrates included Gwo1, which was previously identified to be associated with Argonaute, a core component of RNAi machinery; Pbp1, a homolog of the *Saccharomyces cerevisiae* poly-A-binding protein (Pab1) binding protein, a regulator of mRNA polyadenylation; Puf4, a member of the pumilio-family of RNA binding proteins that was previously shown to be involved in cell wall biosynthesis in *C. neoformans*; and Vts1, a member of the Smaug family of transcriptional regulators that binds to RNA through the sterile alpha motif (SAM) (Mangus *et al*. 1998; Rendl *et al*. 2008; Dumesic *et al*. 2013; Kalem *et al*. 2021). Interestingly, all of the targets of calcineurin localized to P-bodies are involved in RNA binding or processing, with one of them, Gwo1, being part of the RNAi machinery itself (Dumesic *et al*. 2013; Park *et al*. 2016). Based on calcineurin localization in P-bodies and the role of its targets in RNA binding and processing, we explored if calcineurin plays a role in RNAi given that RNAi is executed within P-bodies.

We hypothesized that calcineurin might regulate RNAi through its action on substrates in P-bodies. This was studied by analyzing the possible roles of calcineurin and its P-body localized substrates in MIS (Wang *et al*. 2010). Our results suggest that calcineurin and its substrates contribute to transgene silencing although none are essential for silencing during mitotic growth. We also found that calcineurin plays a role in the reactivation of silenced transgenes. Based on genetic epistasis analysis, calcineurin, and its substrates regulate transgene silencing in different fashions. Genome-wide small RNA sequencing revealed that calcineurin mutations down-regulate small RNA production throughout the genome including from transgenes. Interestingly, *GWO1* deletion abolished small RNA from specific loci in the genome without impacting the majority of RNAi targets throughout the genome. Overall, our studies identify a novel role for calcineurin in RNAi-mediated silencing as well as small RNA production that may contribute to genome defense in *C. neoformans*.

## Materials and Methods

### Strains and media

*C. neoformans* strain JF289**a** served as the reference strain for mitotic-induced silencing (MIS) assays and the congenic H99α strain served as the wild-type isogenic parental for sex-induced silencing (SIS) assays (Perfect *et al*. 1993; Wang *et al*. 2010). Strains were grown in YPD media for all experiments at 30°C for non-selective growth conditions. G418 and/or NAT were added at a final concentration of 200 and 100 µg/ml, respectively, for the selection of transformants. MS media was used for all the mating assays, which were performed as described previously (Sun *et al*. 2019). Random spore dissections were performed after two-three weeks of mating, and the spore germination frequency was scored after five days of dissection. All strains and primers used in this study are listed in Supplementary Table S1 and Supplementary Table S2, respectively.

### Mitotic-induced silencing and unsilencing assays

The frequency for sex-induced and mitotic-induced silencing for the wild-type and various mutants was calculated as described previously (Wang *et al*. 2010). Briefly, for MIS assays, single colonies were obtained for each strain on SD-URA media to ensure expression of *URA5* and incubated at 30°C for two days. For each strain, 3 to 5 single colonies were inoculated in YPD and grown overnight at 30°C per experiment. From the overnight culture, various dilutions were plated on YPD and 5-FOA-containing media. The colonies were counted in the YPD media after 2 days and 5-FOA-containing media after 7 to 10 days and the frequency for mitotic-induced silencing was calculated.

The colonies obtained on the 5-FOA media from the MIS assays were directly resuspended in 100 µl dH_2_O. From this resuspension, 5 µl was directly spotted on the SD-URA media. The colonies exhibiting a robust growth i.e., uniform growth in at least 50% of the spotted area were considered as reverted colonies and the frequency of reversion was calculated.

### PacBio sequencing and genome assembly

Long-read PacBio sequencing was conducted to assemble the complete transgene array for the strain JF289 along with its chromosome-level genome assembly. DNA isolation was performed to enrich the large fragment as described previously (Yadav *et al*. 2020). After a quality check, the DNA was submitted to Duke University Sequencing and Genomic Technologies Core facility for PacBio sequencing. The sequencing reads obtained were then subjected to a de novo genome assembly using Canu 2.0. This resulted in an assembly for all 14 chromosomes, including the full sequence for the transgene array. The genome assembly was then error-corrected (5X) with Pilon using short-read Illumina sequencing reads. The following genome assembly was then employed for the downstream analysis of small RNA reads. The genome annotations were generated by transferring the *C. neoformans* H99 genome annotations onto the JF289 genome based on sequence identity and have not been curated.

### Small RNA preparation and analysis

Single colonies obtained on either 5-FOA or YPD media from the MIS assays were used for small RNA sample preparation and analysis. Small RNA preparation and analysis were performed as described previously (Priest *et al*. 2022). Briefly, colonies were inoculated in 8 ml YPD medium and grown overnight at standard laboratory conditions. Cells were harvested, frozen, and then subjected to lyophilization overnight. 50 to 70 mg of lyophilized material was used for sRNA isolation following the mirVana miRNA Isolation Kit manufacturer’s instructions. sRNA was quantified with a Qubit 3 Fluorometer and submitted for sequencing at the Duke University Sequencing and Genomic Technologies Core facility. sRNA libraries were prepared with a QiaPrep miRNA Library Prep Kit and 1 x 75 bp reads were sequenced on the Illumina NextSeq 500 System.

Post-sequencing, the small RNA reads were processed with cutadapt to remove the adapter sequences, and reads smaller than 14-bp were discarded. The reads obtained after trimming were mapped to the reference JF289 genome using Bowtie2 and resulting SAM files were analyzed for reads length distribution and 5’-nucleotide preference with tools described previously (Priest et al. 2022). For read mapping, we used the default option in Bowtie2 that allows mapping of each read to only one location in the genome based on the mapping quality in order to avoid artificial accumulation of siRNA reads at transposons and repeats. GraphPad Prism was used to calculate percent reads and plot the graphs presented in the figures. Furthermore, SAM files were then converted to BAM files and then to TDF format. The TDF files were visualized in the IGV for their distribution pattern as maps that are presented in figures. The TDF normalization, which multiplies each value by [1,000,000/total read count] for each sample, was employed for visualization in IGV to account for different coverage between samples. The maps were combined with the annotations from the Geneious Prime 2022.1.1 (https://www.geneious.com/) software to overlay the annotation tracks with the read mapping.

Differential expression analysis for the small RNAs was conducted using Geneious Prime. For this purpose, the trimmed reads were mapped to the newly generated JF289 reference genome using Bowtie2, and plug-in DEseq2 (using “parametric” Fit Type) was used to calculate the differential expression of sRNA levels between samples (See details in Supplementary Materials and Methods). The analysis was conducted in two different manners; 1) where only two 5-FOA-specific samples were compared, and 2) where all three samples (two 5-FOA and one YPD) were pooled together. The loci with differential expression are shown in supplementary table S3.

## Results

### Calcineurin deletion results in defective transgene silencing and reactivation

To probe the possible role of calcineurin in RNAi, we employed a previously characterized system to analyze transgene array silencing in *C. neoformans* (Wang *et al*. 2010). The transgene array includes the *URA5* gene arranged in a tandem fashion as a repeat cluster where the expression of the *URA5* gene can be suppressed by RNAi-mediated small RNA generation. The silencing of *URA5* within the transgene was initially discovered during sexual reproduction where it happens in 50% of the progeny that inherit the transgene and requires an active RNAi machinery (Wang *et al*. 2010). This process was defined as sex-induced silencing (SIS) and was shown to act as a defensive mechanism for the *C. neoformans* genome. An analysis of silencing during mitotic growth found transgene silencing, albeit at a much lower frequency of approximately 1/1000, and is defined as mitotic-induced silencing (MIS). Because calcineurin mutants fail to undergo sexual reproduction, we measured the frequency of MIS in the wild-type and isogenic mutant strains. We generated two independent calcineurin A (Cna1) and calcineurin B (Cnb1) mutants in the reporter JF289 strain background that bears the *URA5* transgene array, and these mutant strains were then subjected to MIS RNAi silencing assays (Figure 1A-B). Specifically, isolated single colonies were grown in rich media and then plated to synthetic defined (SD) media containing 5-Fluoroorotic Acid (5-FOA). 5-FOA is metabolized into a toxin in cells expressing *URA5*, and thus only colonies that do not express *URA5* or harbor a *ura5* mutation grow on 5-FOA-containing medium. With this assay, calcineurin mutant strains exhibited an approximate two-fold reduction in transgene RNAi silencing as compared to the wildtype parent strain JF289, suggesting a modest but demonstratable role for calcineurin in this process (Figure 1C-D). To further assess this, we performed MIS assays after treating JF289 cells with the calcineurin inhibitor FK506 in an overnight culture and then plated cells on 5-FOA media containing FK506. This experiment revealed that prolonged calcineurin inhibition had a similar impact on the MIS frequency (Figure 1D). As expected, a known RNAi mutant (*rdp1*Δ) did not produce any 5-FOA resistant colonies in MIS assays. Calcineurin mutation or inhibition confers growth defect at temperatures higher than 25°C, we directly tested whether such a growth defect could lead to MIS reduction in these mutants. First, we measured the doubling time of these mutants at 30°C which showed that both the *cna1*Δ and the *cnb1*Δ mutants exhibited slower growth growing 1.3-1.5 times slower than the wild-type (Table S4). We also measured the growth rate in SD media (which is supplemented with 5-FOA for assessing MIS frequency) and found a similar reduction in growth rate for the mutants. Next, we measured the viability of these strains in these two media conditions and found a similar loss in viability in the mutants in both YPD and SD media (Figure S1A). Combined together, these results show that the mutants are similar in both nutrient-rich YPD and nutrient-limiting SD media. During the MIS frequency calculation, we always factor in the number of viable cells plated by counting colonies on the YPD plate (Figure 1B), which accounts for the potential loss of viability in these mutants, negating any impact on MIS frequency as a result of viability loss. Next, we measured MIS frequency in the wild-type and two calcineurin mutants after growth for 18 hours and 40 hours from the same culture to determine the direct impact of the number of cell divisions. We found that the MIS frequency was unchanged for all three strains over time indicating the number of cell divisions had little to no impact on transgene silencing in these strains (Figure S1B). Considering the epigenetic nature of this phenomenon, where a given cell could switch between silenced and unsilenced states at any point of time irrespective of the number of divisions, this might be a predictable finding. Based on these results, we conclude that MIS reduction observed in calcineurin mutants is not a result of cell growth defects.

**Figure 1.**
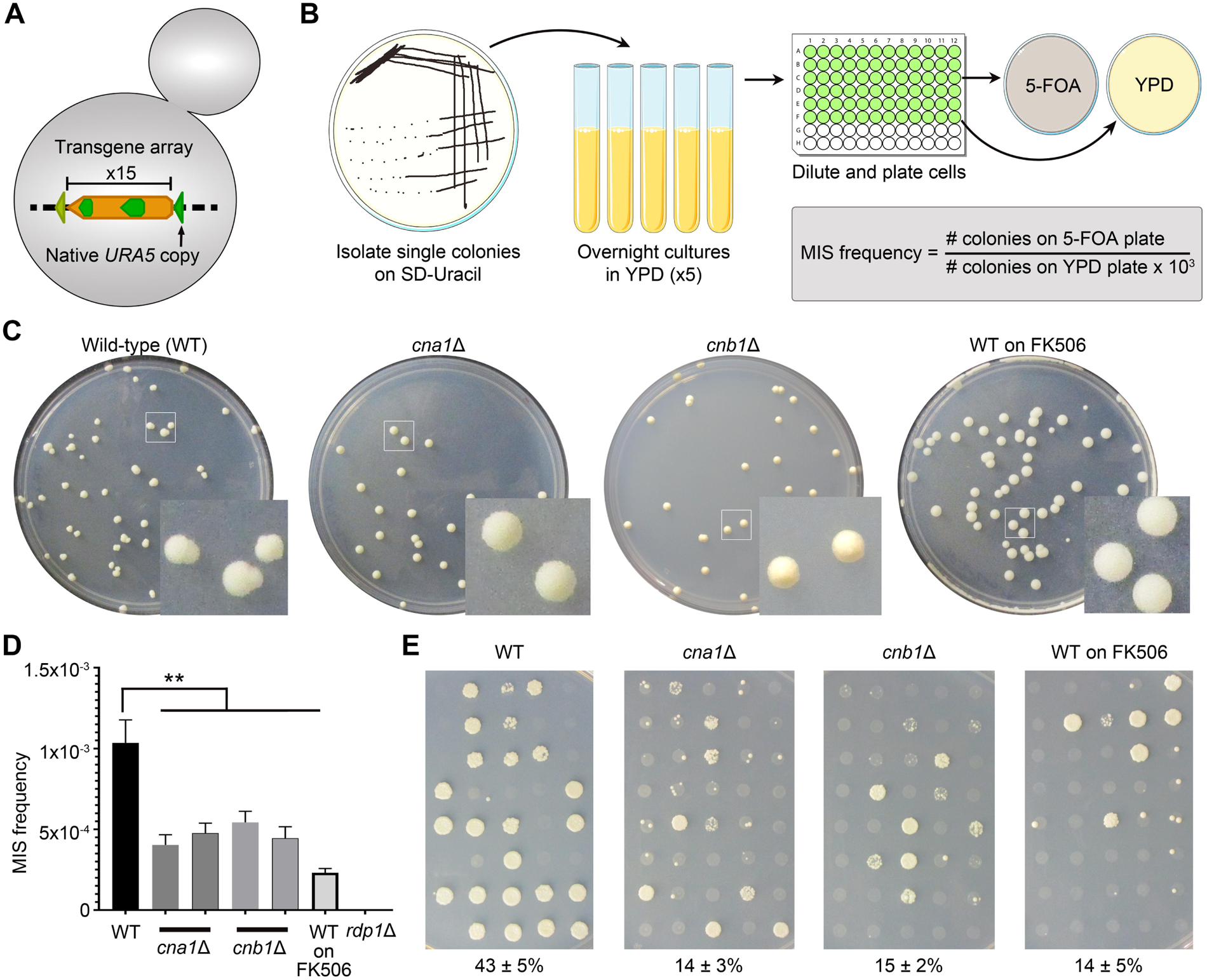
Calcineurin regulates the onset and reversion of transgene silencing. **(A)** A cartoon depicting the *URA5* transgene array in the reporter strain JF289. **(B)** A schematic showing the workflow to determine the mitotic-induced silencing (MIS) frequency with the reporter strain. **(C)** Plate photos showing the difference in colony morphology for the wild-type JF289, *cna1*Δ, *cnb1*Δ, and JF289 plated on FK506 containing media when an approximately equal number of colonies were obtained in each case after 10 days of incubation. **(D)** A graph showing the MIS frequency of calcineurin mutants as compared to the JF289 strain. P-value (**) < 0.01. Error bars represent the standard error of the mean (SEM). **(E)** Images of petri-dishes showing the growth of colonies on media lacking uracil (SD-URA) directly from 5-FOA-containing media during MIS assays. The numbers below represent the average percent of colonies growing (Mean ± SEM) from three independent experiments.

During the MIS assays, we observed that the colony morphology for calcineurin mutants on 5-FOA media as well as for the wild-type cells treated with FK506 differed from the wild-type parent without any drug treatment (Figure 1C). Notably, the majority of colonies (>70%) obtained in the calcineurin mutants or upon calcineurin inhibition were homogeneous with smooth edges, in stark contrast to those from the wild-type strain where >80% of the colonies were sectored with rough peripheries. We hypothesized that this might be attributable to a difference in the rate at which cells within the colonies reverted to *URA5* expression and that calcineurin mutation might interfere with the reversion of RNAi-mediated transgene silencing. To test this, we scored >100 colonies from 5-FOA media for their ability to grow on a defined medium lacking uracil (SD-URA) that only allows the growth of cells that express *URA5*. Colonies that exhibited a uniform growth in more than 50% of the spot were counted as reverted or unsilenced whereas the rest were counted as silenced. These experiments revealed that a significantly higher number of JF289 colonies could directly grow on SD-URA, as compared to calcineurin mutants or JF289 cells growing in the presence of FK506 (Figure 1E). This result suggests that calcineurin plays a role in controlling the frequency at which a silenced transgene reverts and thereby escapes RNAi silencing. We also tested the MIS reversion frequency for calcineurin mutants from the time-point experiment and found that colonies from both 18 hours and 40 hours time points reverted at a similar rate (Figure S1B). Taken together, we conclude that calcineurin mutants are not only modestly defective in silencing the *URA5* gene, but also exhibit a reduced rate of unsilencing, thus contributing to prolonging the state of RNAi-mediated transgene silencing.

### Calcineurin substrates in P-bodies play different roles in silencing

To understand how calcineurin regulates RNAi silencing, we hypothesized that calcineurin targets/substrates in P-bodies might be involved in this process (Park *et al*. 2016). To test this, we generated and tested two independent mutants of known calcineurin substrates that localize to P-bodies and bind to RNA (Gwo1, Pbp1, Puf4, and Vts1) in the transgene array strain background (JF289) and determined the MIS frequency for these mutants. Interestingly, *gwo1*Δ mutations led to an approximately five-fold reduction whereas *puf4*Δ mutations led to an approximately three-fold increase in MIS (Figure 2A). While no significant difference in the MIS frequency was observed for the *pbp1*Δ mutants, we did observe more heterogeneity in the shape and sizes of the colonies, suggesting that silencing might be delayed in the *pbp1*Δ mutant without an alteration in the overall rate of silencing (Figure 2B). Finally, the *vts1*Δ mutant did not exhibit any difference in the MIS frequency. Unlike the calcineurin mutants, none of these mutants exhibited a growth defect (Table S4) ruling this out as a possible cause of the MIS defect in both the *gwo1*Δ and the *puf4*Δ mutants. Next, we tested the frequency of transgene silencing reversion for the mutants of the four P-body components by directly transferring the colonies from 5-FOA media to SD-URA media. These experiments showed that *puf4*Δ mutations completely abolished the MIS reversion as none of the colonies exhibited uniform growth on uracil lacking media. Deletion of the other three genes did not have a significant impact on transgene reactivation with the frequency ranging between 30 to 39% (Figure 2C).

**Figure 2.**
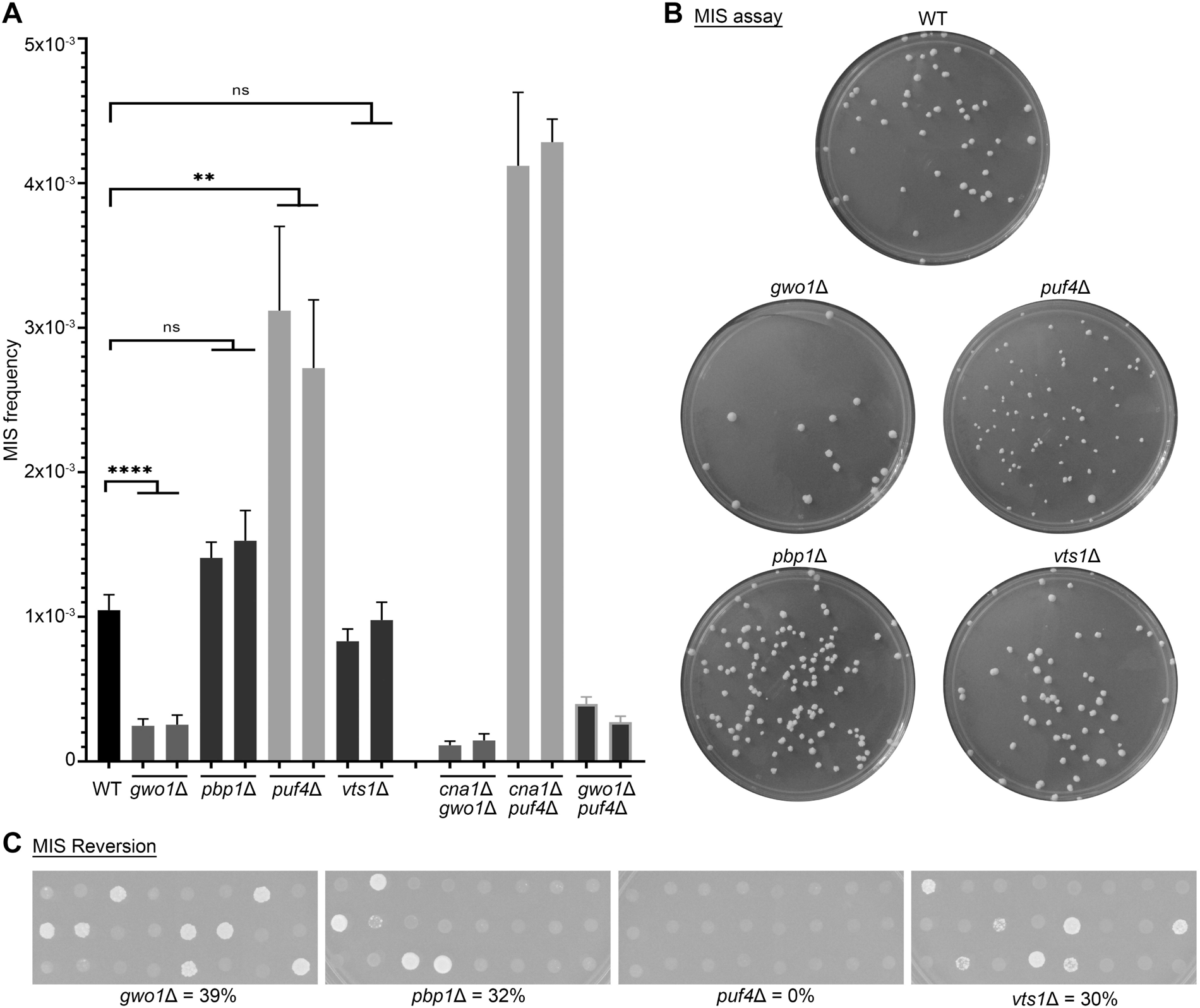
Calcineurin substrates play diverged roles during transgene silencing. **(A)** A graph showing the MIS frequency of mutants for known calcineurin targets (Gwo1, Pbp1, Puf4, and Vts1) as well as double deletion mutants of genes that showed an effect on MIS (p-values: ** < 0.01, **** < 0.0001). The double deletion mutants were not found to be significant as compared to their respective mutants (p-value >0.05). **(B)** Photographs of petri-dishes showing the colony number variation in different mutants, as compared to JF289 after 10 days of incubation when an estimated equal number of cells were plated in each case. **(C)** Images showing the growth of individual colonies on SD-URA media from 5-FOA containing media for MIS reversion assays. The numbers represent the reversion rates for each mutant.

Among these targets, *gwo1*Δ and *vts1*Δ mutants can successfully undergo sexual reproduction and produce spores whereas *puf4*Δ and *pbp1*Δ mutations impair sexual reproduction. This allowed us to determine the RNAi silencing frequency during sexual reproduction (SIS) for the *gwo1*Δ and v*ts1*Δ mutants. For this purpose, we generated mutants for these genes in a congenic strain of opposite mating-type, H99α, and crossed them with respective mutants in the transgene reporter strain JF289**a**. The basidiospores produced from these crosses were randomly dissected by micromanipulation to obtain viable progeny on the non-selective media (YPD). The germinated spores were then genotyped for the presence of the *URA5* transgene in their genome and in parallel assayed for their growth on 5-FOA-containing media. SIS frequency was then calculated as the number of progeny growing on 5-FOA-containing media out of the total number of progeny that inherited the transgene array. Previous studies have established that a wild-type cross results in *URA5* silencing in approximately 50% of the progeny (Wang *et al*. 2010; Feretzaki *et al*. 2016) and it occurred at ∼45% in our experiments (Table 1). Interestingly, *gwo1*Δ mutations led to a complete abolishment of SIS in the H99 *gwo1*Δ x JF289 *gwo1*Δ cross and none of the analyzed progeny grew on 5-FOA. By comparison, a 30% reduction (SIS frequency ∼33%) in SIS was observed in a cross between H99 *vts1*Δ between JF289 *vts1*Δ strains. These results indicate that the impact of Gwo1, and probablyVts1, on RNAi-mediated silencing is considerably more evident during sexual reproduction. Overall, these results suggest that calcineurin targets in P-bodies are involved in RNAi-mediated silencing but fulfill different roles suggesting the presence of multiple pathways controlling this process.

**Table 1.** SIS frequency for JF289, *vts1*Δ, and gwo1Δ mutant crosses.

| Cross | Viable progeny obtained | Progeny inheriting transgene array |  | SIS frequency |
| --- | --- | --- | --- | --- |
|  |  | Total progeny | Growth on 5-FOA |  |
| H99α x JF289a | 71 | 36 | 16 | 44.4% |
| H99α *vtl1Δ* x JF289a *vtl1Δ* | 59 | 27 | 9 | 33.3% |
| H99α *gwo1Δ* x JF289a *gwo1Δ* | 92 | 49 | 0 | 0% |

### Calcineurin and its targets impact RNAi-mediated silencing through multiple pathways

Previously, studies showed that Gwo1, Pbp1, Puf4, and Vts1 localize more prominently to P-bodies in response to heat stress (Park *et al*. 2016). Here, we first explored if calcineurin controls the localization of these targets and thus impacts RNAi-mediated silencing. To study this, we performed co-localization of these calcineurin targets at 37°C in the presence of the calcineurin inhibitor, FK506. Interestingly, these experiments revealed that the localization pattern of all four proteins is independent of calcineurin activity and they co-localize with a P-body marker, Dcp1, irrespective of calcineurin activity (Figure S2). Similar to a previous study (Kozubowski *et al*. 2011), we also noted that calcineurin activity is not required for its own localization to P-bodies (Figure S2 – top row). These results suggest that the localization of calcineurin and its targets to P-bodies may be driven by other factors such as structural determinants.

To further understand the connection between calcineurin and its targets for their roles in RNAi-mediated silencing, we generated double deletion mutants lacking Cna1 in combination with loss of Puf4 or Gwo1 because these two targets exhibited significant differences in MIS frequency. Surprisingly, both *cna1*Δ *gwo1*Δ and *cna1*Δ *puf4*Δ mutants exhibited MIS frequencies similar to *gwo1*Δ and *puf4*Δ single mutants, respectively (Figure 2A). These results indicate a more dominant role for both Puf4 and Gwo1 than Cna1 in MIS silencing. It is also possible that calcineurin-mediated dephosphorylation may not be required for the role of Gwo1 or Puf4 in RNAi-mediated silencing. Furthermore, calcineurin might have additional substrates in P-bodies that regulate MIS in different ways negating the effects of calcineurin in these double mutants. Because Gwo1 and Puf4 regulate MIS in opposite directions, we also generated a *gwo1*Δ *puf4*Δ double mutant and determined the MIS frequency. We attempted to obtain this double mutant strain by crossing the strains lacking either one of the genes in the parents and analyzing the progeny by PCR analysis. We failed to obtain a *gwo1*Δ *puf4*Δ double mutant from H99 *gwo1*Δ x JF289 *puf4*Δ (0 out of 73 viable progeny) suggesting the double mutant might have a fitness defect. *GWO1* is located on chromosome 1 whereas *PUF4* is present on chromosome 3, ruling out linkage as an explanation for this observation. Interestingly, we recovered 11 progeny that were *gwo1*Δ *puf4*Δ double mutants from a reciprocal cross H99 *puf4*Δ x JF289 *gwo1*Δ (11 out of 80), albeit at a lower frequency than expected (25%). The MIS assays with the *gwo1*Δ *puf4*Δ double mutant revealed the MIS frequency at the same rate as that of the *gwo1*Δ single mutant suggesting that Gwo1 plays a more dominant role in transgene silencing than Puf4 (Figure 2A). Overall, these studies show that calcineurin and its known targets impact RNAi-mediated transgene silencing through multiple pathways that may or may not be connected with each other.

### Calcineurin mutants produce less RNAi-mediated small RNA from the transgene

A previous study showed that transgene silencing is mediated by RNAi-mediated small-interfering RNAs (siRNAs) (Wang *et al*. 2010). We therefore assessed if calcineurin and its targets impact siRNA production from the transgene array. We first conducted long-read PacBio sequencing to assemble the genome of the transgene array containing reporter strain, JF289. The genome assembly revealed 15 tandem copies of the transgene integrated next to the endogenous *URA5* gene. After assembling the complete transgene array, we conducted small RNA sequencing from the wild-type transgene array strain JF289 as well as mutant strains.

We selected two 5-FOA resistant colonies (transgene silenced) obtained from the MIS assays and one YPD colony (transgene expressed) from the wild-type (JF289) as well as mutants that exhibited differences in MIS silencing (*cna1*Δ, *cnb1*Δ, *gwo1*Δ, and *puf4*Δ strains). Following sRNA-sequencing, the reads were processed to trim adapters followed by mapping to the reference genome sequence. This mapped small RNA population was first analyzed the small RNA population for RNAi-mediated signatures such as RNA length and 5’ nucleotide preference. This analysis revealed that all strains are proficient in RNAi-mediated siRNA production and exhibited a peak at 21-23 nucleotides with a 5’-Uracil preference, both signatures of canonical RNAi processing (Figure 3). Interestingly, the *cna1*Δ and *cnb1*Δ mutants exhibited a significant 30-40% reduction in this population of siRNA compared to the wild-type and other two mutants (Table 2). These results confirm the role of calcineurin in small RNA production and provide additional evidence for its role in transgene silencing.

**Figure 3.**
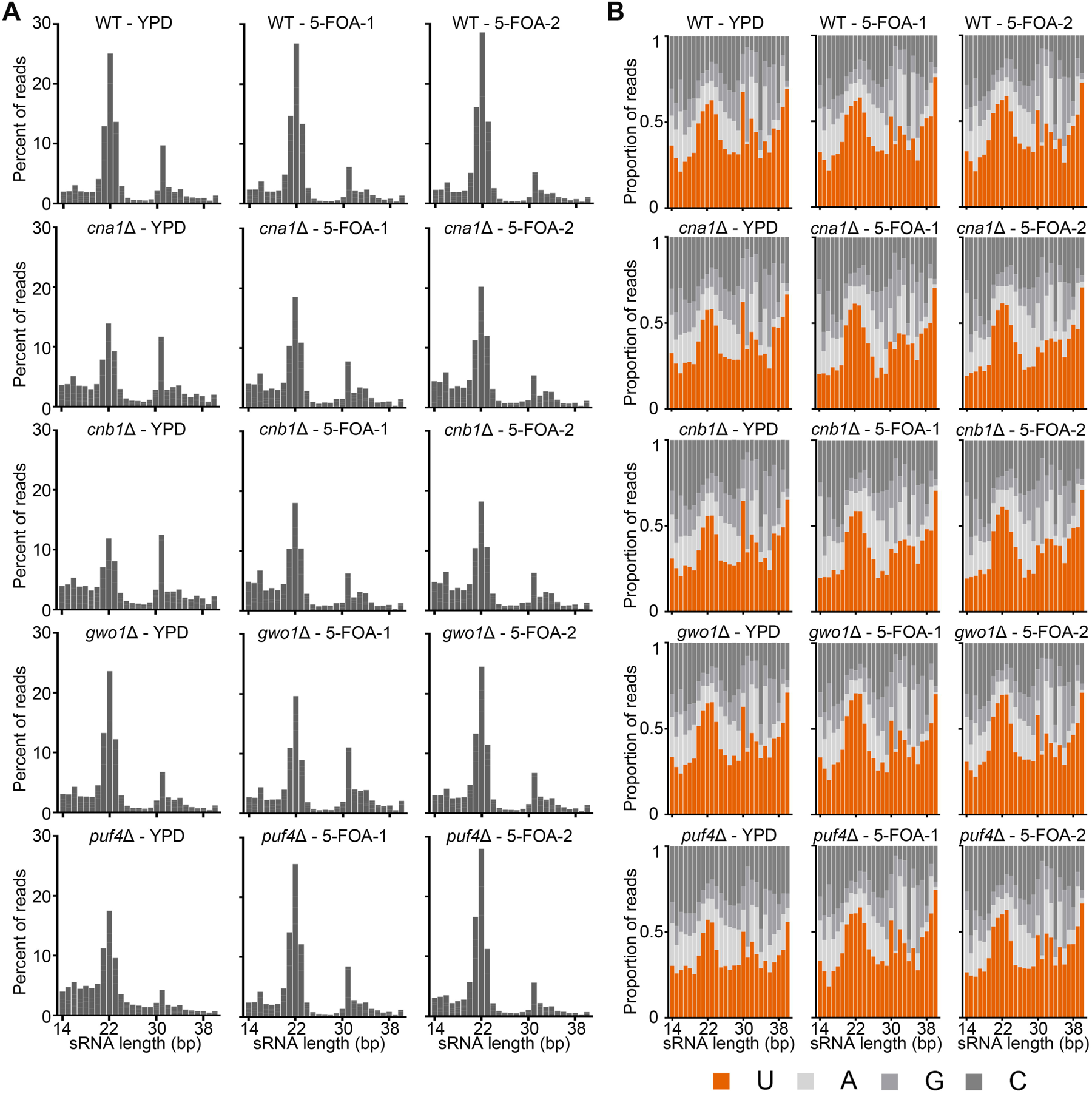
Calcineurin and its substrates are not essential for RNAi-mediated siRNA production. Graphs showing the (A) length distribution and (B) 5’ nucleotide preference of small RNAs in the wild-type (JF289) and mutant strains. The isolates analyzed for the small RNA preparation were from both non-selective (YPD) and selective (5-FOA) conditions for the MIS assays.

**Table 2.** siRNA population levels in the wild-type and various mutants.

| Condition | siRNA length | Percent of reads in siRNA population |  |  |  |  |
| --- | --- | --- | --- | --- | --- | --- |
|  |  | Wild-type | *cna1Δ* | *cnb1Δ* | *gwo1Δ* | *puf4Δ* |
| **YPD** | 21-nt | 7.30 | 4.08 | 3.48 | 8.06 | 5.57 |
|  | 22-nt | 15.15 | 8.02 | 6.66 | 15.30 | 10.02 |
|  | 23-nt | 8.58 | 5.40 | 4.55 | 8.02 | 5.34 |
|  | <u>Total</u> | 31.03 | 17.50 | 14.69 | 31.38 | 20.93 |
| **5-FOA-1** | 21-nt | 8.68 | 6.01 | 5.78 | 7.33 | 8.56 |
|  | 22-nt | 16.57 | 11.39 | 10.62 | 13.89 | 15.54 |
|  | 23-nt | 8.54 | 6.59 | 6.13 | 6.33 | 7.82 |
|  | <u>Total</u> | 33.79 | 23.99 | 22.53 | 27.55 | 31.92 |
| **5-FOA-2** | 21-nt | 9.69 | 6.52 | 5.94 | 8.68 | 9.65 |
|  | 22-nt | 17.85 | 12.47 | 11.19 | 17.14 | 16.88 |
|  | 23-nt | 8.86 | 7.29 | 6.32 | 8.04 | 7.07 |
|  | <u>Total</u> | 36.40 | 26.28 | 23.45 | 33.86 | 33.60 |
| Mean ± SEM of total siRNA/condition |  | 33.74 ±1.55 | 22.59 ±2.63 | 20.22 ±2.78 | 30.93 ±1.84 | 28.81 ±3.97 |
| Significant; p-value (Compared to wild-type) |  |  | Yes; 0.011 | Yes; 0.012 | No; 0.278 | No; 0.199 |

Next, we mapped the small RNA reads to the JF289 genome and first analyzed the reads mapping specifically to the transgene array (Figure 4). In accordance with the previous report (Wang *et al*. 2010), we identified siRNA only against the *URA5* component of the transgene, and no siRNA peaks were identified against the rest of the transgene including the *SXI2***a** gene. More specifically, this siRNA population against the *URA5* gene was present only in the cultures grown from 5-FOA colonies (*URA5* silenced), but not in the YPD colonies cultures (*URA5* expressed) (Figure 4). The presence of RNAi-mediated siRNA population in these 5-FOA colonies is expected because they are required for suppression of *URA5* expression, whereas YPD colonies have not been selected for *URA5* silencing in accord with the lack of siRNAs. Both *cna1*Δ and *cnb1*Δ mutants exhibited fewer siRNAs against *URA5* as compared to the wild-type (Figure 4), in accordance with the overall reduction in RNAi-mediated siRNA population in these mutants as described above. In our analysis, we mapped each read to only one location in the genome and as a result all reads generated from the transgene locus are uniformly distributed across all copies of *URA5*. It is possible that some *URA5* copies might produce more siRNA than others but if it occurs it cannot be inferred from this analysis.

**Figure 4.**
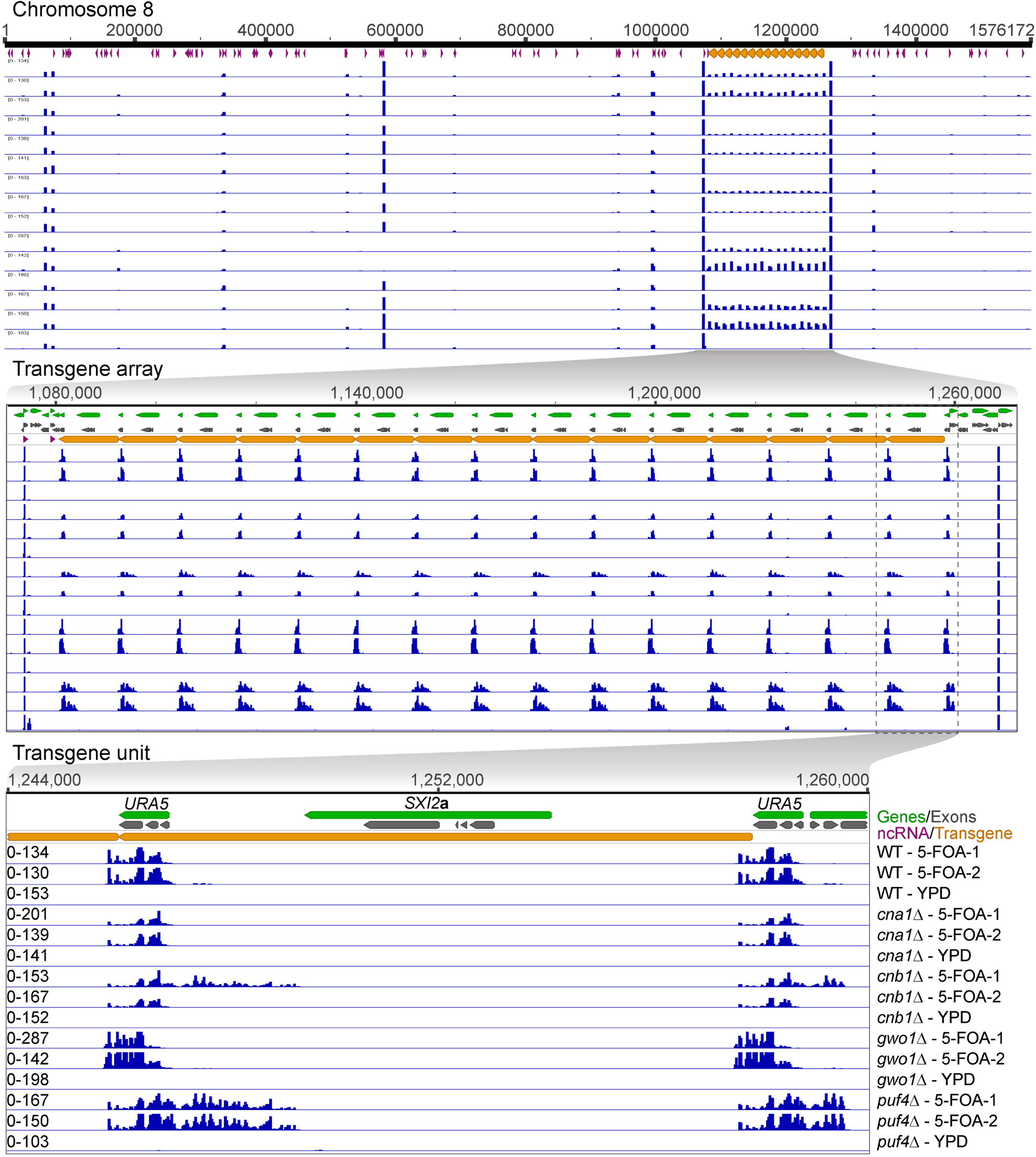
Calcineurin mutants produce less siRNA against the transgene array. A series of genomic location maps showing the status of siRNA against the transgene array is presented with the top panel showing the full chromosome 8 on which the transgene array is integrated. The middle panel shows a zoomed view of the entire transgene array and mapping of siRNA from the wild-type as well as various mutants. The lower panel depicts a single unit of transgene array including the location of *URA5* and *SXI2***a** genes. *URA5* outside of the transgene array represents the endogenous copy of *URA5* that was targeted for transgene insertion. The mapped siRNA data is presented in the same order across all three panels and is labeled in the lower panel. The siRNA mapping revealed a lower level of siRNA in both *cna1*Δ and *cnb1*Δ mutants whereas a difference in siRNA distribution was observed in *gwo1*Δ and *puf4*Δ mutants.

Further analysis revealed mutant-specific behavior of the siRNA population where both *gwo1*Δ and *puf4*Δ mutants exhibited a change in siRNA distribution. *gwo1*Δ mutant isolates produced more siRNA against the 3’ end of the *URA5* gene and less against the 5’ end as compared to the wild-type transgene array strain JF289. On the other hand, *puf4*Δ mutant isolates generated similar levels of siRNA against *URA5* to that of wild-type but additionally produced siRNAs from the upstream region of the *URA5* gene spreading into the neighboring gene. A similar pattern of siRNA was also observed in one of the *cnb1*Δ mutant 5-FOA isolates, albeit at a lower level. These results reveal that calcineurin, Gwo1, and Puf4 regulate the siRNA population in different manners affecting either the level or the profile of siRNAs against the *URA5* gene.

### Genome-wide siRNA production is altered in calcineurin and gwo1***Δ*** mutants

After analyzing the transgene-specific small RNAs, we next assessed whether calcineurin mutants also affect small RNA production on a genome-wide level. Analysis of the small RNA levels throughout the genome revealed that both the *cna1*Δ and *cnb1*Δ mutants produced fewer siRNAs (Figure 5, S3 and Table 2). To quantify this, a differential expression analysis was performed which revealed that both the *cna1*Δ and *cnb1*Δ mutants exhibited a change in the production of siRNAs against several loci with the *cnb1*Δ mutation affecting at slightly more loci (Table S3). This comparative analysis also showed that the *gwo1*Δ mutant also impacted a large number of loci whereas the *puf4*Δ mutant altered siRNA levels from only a few loci, suggesting a more dominant role for Gwo1 in RNAi in accord with results from the MIS assays.

**Figure 5.**
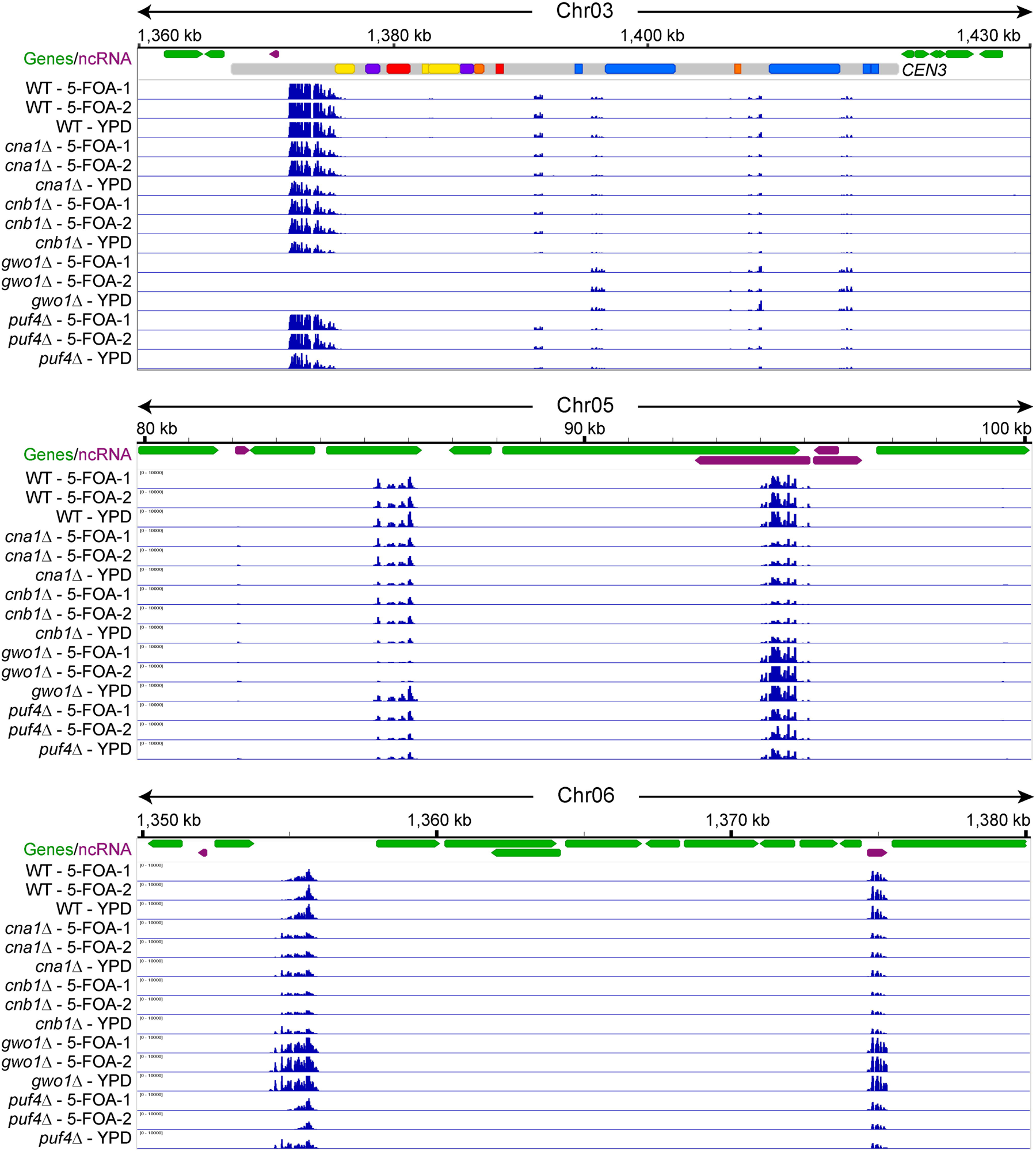
Calcineurin and Gwo1 regulate genome-wide siRNA production differently. Snapshots of various genomic regions showing small RNA reads mapping for various strains including calcineurin mutants. The top panel depicts a centromere from chromosome 3, whereas other panels present two other genomic loci. These regions show similarities and differences in the siRNA distribution profiles for the wild-type and mutant strains. Also see Figure S3.

Most genomic regions exhibited mutant-specific differential expression where the siRNA population from certain loci showed changes across all samples, two from 5-FOA, and one from YPD. However, some loci produced different levels of siRNA depending on the condition for the same mutant. For example, siRNA production from a region in centromere 3 and an intergenic region on chromosome 6 was different between YPD and 5-FOA samples in the *puf4*Δ mutant (Figure 5, top and bottom panels). A more striking example was observed for the *gwo1*Δ mutant which showed no siRNA from a locus on chromosome 8 in the 5-FOA condition and abundant siRNA production from the same loci in the YPD condition (Figure S3, left side, middle panel). While these regions do not exhibit any similarity to the *URA5* gene, whose expression 5-FOA counter-selects, it is important to note that 5-FOA-containing media provides a stressful growth condition and these siRNA changes observed may be due to this additional stress resulting in epi-alleles at these loci.

The genome-wide analysis also revealed that the *gwo1*Δ deletion affects siRNA production from certain loci more dramatically than others. For several loci, *gwo1*Δ deletion caused a complete abolishment of siRNA production (Figure 5 and S3), whereas for many others this led to an increase in siRNA production (compare various loci shown in the upper two panels of Figure 5). This particular pattern was observed for multiple loci throughout the genome showing that the *gwo1*Δ mutation has a significant impact on siRNA production. Overall, siRNA sequencing analysis revealed that calcineurin and Gwo1 are major factors in the global regulation of small RNAs in *C. neoformans*.

## Discussion

In this study, we showed that calcineurin contributes to RNAi-mediated silencing as well as the reactivation of a transgene array. Our results suggest that this role of calcineurin may be exerted both through and independent of its known targets in P-bodies, and requires interactions with the RNAi machinery (Figure 6). Interestingly, one of the known substrates, Gwo1, has been identified in an earlier study (Dumesic *et al*. 2013) as a component of the RNAi machinery via its binding to Argonaute (Ago1), a core component of the RNAi machinery, and notably showed the largest impact on transgene silencing both during MIS and SIS in our assays. Interestingly, Dumesic et al had concluded that Gwo1 is not required for siRNA biogenesis based on the analysis of three genomic loci. Contrary to that, our analysis identified that loss of Gwo1 has a locus-specific impact and is required for siRNA biogenesis from certain genomic loci but not others, unlike conventional RNAi proteins. We also identified that deletion of Puf4 results in enhanced silencing by approximately three-fold and this is the first gene whose mutation has been identified to enhance silencing. Puf4 belongs to the Pumilio and Fem-3 (PUF) family of mRNA binding factors and has been shown to bind to several RNAs in *C. neoformans* (Kalem *et al*. 2021; Kalem *et al*. 2022). RNA interactions of Puf4 involve genes required for cell wall synthesis, drug resistance, as well as regulating gene expression. More importantly, Puf4 immuno-precipitation also revealed a direct interaction with Ago1 (Kalem *et al*. 2022). It is possible that both Gwo1 and Puf4 controls siRNA loading on Ago1 and future studies would explore siRNA cargo of Ago1 in the absence of these two proteins. Our results also indicate a more global role for calcineurin than either Gwo1 or Puf4 indicating that calcineurin might be regulating siRNA production and transgene silencing independently of these two substrates. Such a global impact might involve either a direct interaction of calcineurin with the core RNAi machinery or an indirect effect through one of its uncharacterized substrates that plays a major role in RNAi function. Delineating these direct and indirect effects of calcineurin on RNAi-mediated transgene silencing and siRNA production will be crucial to better understanding the impact of calcineurin signaling in genome defense mechanisms.

**Figure 6.**
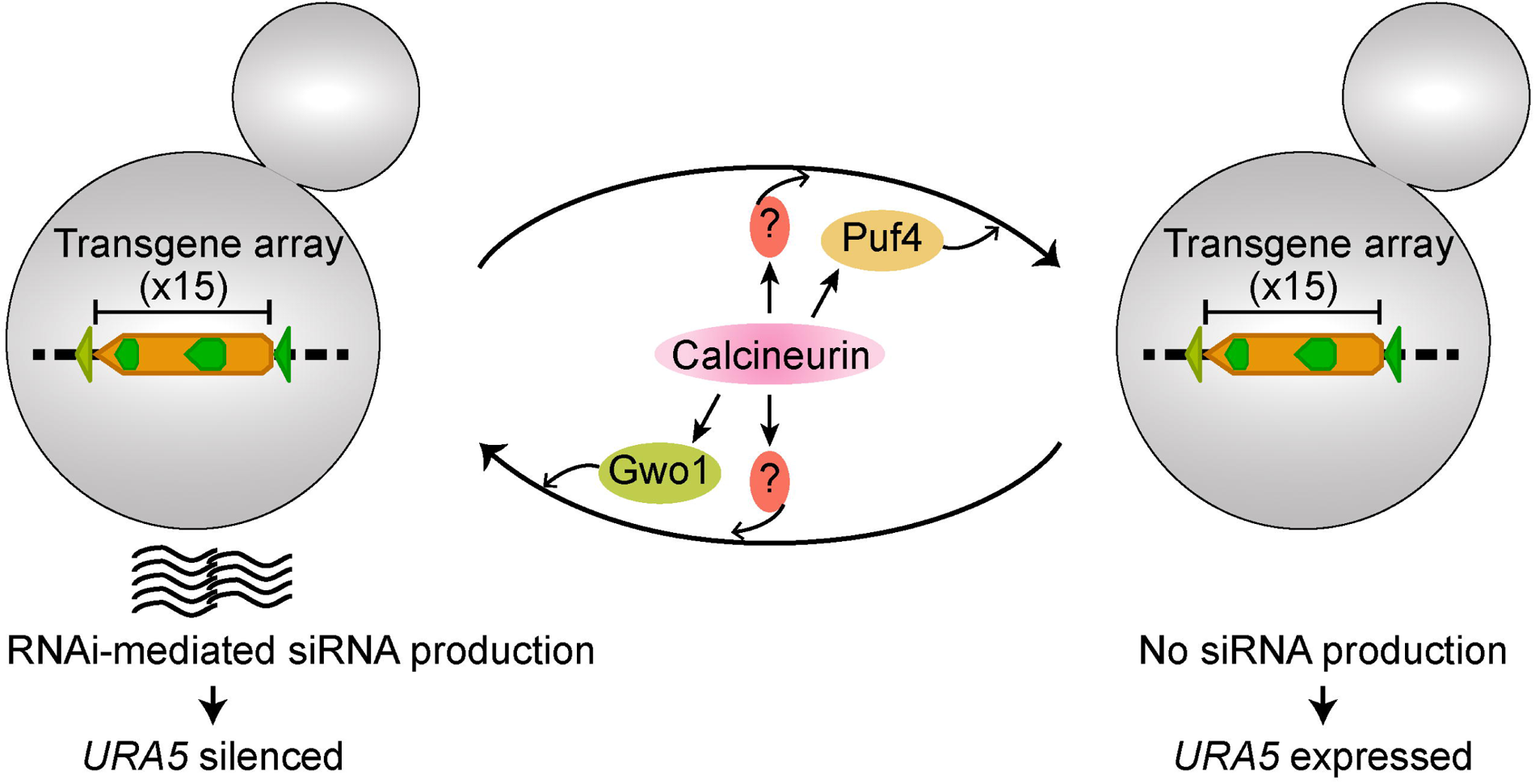
A model showing the role of calcineurin, and its known substrates, Gwo1 and Puf4, in regulating RNAi-mediated small RNA production.

One of the main findings of this study is the demonstration of a role for calcineurin in reducing both silencing and un-silencing of the transgene array, revealing that calcineurin maintains the flux between the silenced and expressed states. Three of four calcineurin substrates analyzed in this study (Gwo1, Pbp1, and Vts1) do not impact the un-silencing of the transgene array whereas Puf4 deletion resulted in the complete abolishment of transgene reactivation. This result suggests that calcineurin could impact the transcription status of cells in response to stress conditions in a previously unknown manner. A recent study in plants identified a link between calcium signaling and antiviral RNAi defense (Wang *et al*. 2021). This study showed that calcium signaling activated calmodulin-binding transcription activator-3 (CAMTA-3) to drive expression changes of key RNAi machinery components in plants. In our study, we show a direct link between RNAi and calcineurin, a phosphatase that is also activated by calcium and calmodulin. Whether this might be conserved in animals is unknown. Combined, these studies indicate that calcium signaling might have a broader role in genome defense mechanisms such as RNAi and might act through different downstream components in different organisms.

Previous studies revealed a large group of genes differently regulated at 37°C in the absence of Cna1 independent of Crz1, a transcription factor that is activated by calcineurin (Chow *et al*. 2017). How calcineurin governs the mRNA abundance of these genes independent of Crz1 is not well understood but might involve post-transcriptional regulation in P-bodies. First, calcineurin activity may be essential for the processing of certain mRNAs, and in its absence, those mRNAs might be either degraded or stored for longer times resulting in an alteration in RNA levels. A second possible mechanism of calcineurin regulation might be through its role in RNAi where calcineurin might be responsible for degrading certain mRNAs via RNAi at high temperatures. The third possible role of calcineurin in regulating RNA metabolism might be through a direct role in the nucleus. Calcineurin localizes to the nucleus in mammalian cells, and this localization varies among different calcineurin isoforms (Shibasaki *et al*. 1996; Jabr *et al*. 2007). While calcineurin has not as yet been reported to localize to the nucleus in *C. neoformans*, it cannot be excluded as a possible mechanism of action. Lastly, it is also possible that calcineurin regulates other transcription factors in addition to Crz1 that drive the expression of a different set of genes. Our experiments employing small RNA sequencing revealed that calcineurin has a significant impact on siRNA production from hundreds of genomic loci providing evidence for the second proposed hypothesis above. However, more than 30% of these affected loci are non-coding RNAs, and siRNA changes could account for only 29 genes whose expression is differentially regulated in the calcineurin mutant based on RNA-sequencing, suggesting that calcineurin’s role on RNA metabolism could be through multiple independent regulatory processes. Additionally, it is possible that the impact of calcineurin on siRNA production is more enhanced at 37°C and could account for a larger number of genes whose expression is altered upon heat stress. No study has analyzed siRNA levels at different temperatures making it a subject of investigation for future studies to better understand the roles of the RNAi machinery in heat stress responses in *C. neoformans*.

RNAi-mediated silencing is known for its roles in genome integrity and defense mechanisms (Obbard *et al*. 2009; Dumesic *et al*. 2013; Yadav *et al*. 2018; Priest *et al*. 2022). A role for calcineurin in RNAi regulation suggests that calcineurin might also be contributing to genome integrity in ways that have not been defined. This is further strengthened by recent studies in humans where calcineurin roles at nuclear pore complexes and centrosomes were discovered (Wigington *et al*. 2020; Tsekitsidou *et al*. 2023). In *Schizosaccharomyces pombe*, RNAi is required for centromere function and if calcineurin has a similar role in regulating RNAi in this fission yeast, it might be directly contributing to genome maintenance and nuclear division (Volpe *et al*. 2003; Folco *et al*. 2008). Similarly, RNAi regulates centromere structure in *C. neoformans* (Yadav *et al*. 2018), and whether calcineurin also plays a role will be of interest to study. Whether calcineurin has broadly conserved or species-specific roles in genome integrity in other organisms will be an interesting avenue to pursue, especially given that no clear link has been established between calcineurin and genome defense mechanisms.

Our results also suggest that processes like RNAi can be affected by environmental stimuli through signaling pathways such as calcineurin. This not only expands the repertoire of factors that affect genome defense mechanisms but also provides important insights on the mechanisms of RNAi-associated processes. For example, epimutations that drive drug resistance in some fungi might have an origin in stress-induced signaling that is initiated in response to infection (Calo *et al*. 2014; Chang *et al*. 2019; Chang and Heitman 2019). In *C. neoformans*, transgene silencing occurs at a much higher rate during meiosis than in mitosis (Wang *et al*. 2010). While it has been shown that RNAi is more active during meiosis, what triggers this change remains to be identified. It is possible that environmental stimuli play an important role in this switch and signaling pathways such as calcineurin may play a role. Calcineurin in *C. neoformans* is not required for vegetative growth at 25°C and cells lacking calcineurin behave like the wild-type at this temperature. However, when calcineurin mutant cells are subjected to mating that also occurs at 25°C, they fail to produce filaments revealing the impact of calcineurin in response to mating-inducing cues specifically (Cruz *et al*. 2001). Previous studies have also identified that the RNAi machinery is activated during the early stages of mating itself, possibly in response to the same environmental stimuli that triggers the calcineurin pathway (Wang *et al*. 2010). Whether there is a direct connection between calcineurin and RNAi during mating and later stages of sexual reproduction remains to be explored.

Overall, our study opens avenues for exploration with respect to the roles of calcineurin in gene expression, maintaining genome integrity, as well as linking environmental stimuli with processes such as RNAi. In this study, we provide a connection between calcineurin signaling and RNAi-mediated silencing that cannot be explained entirely through known substrates of calcineurin. Thus, a better understanding of calcineurin signaling is required to fully understand its roles in *C. neoformans* biology, specifically in stress-response pathways.

## Data Availability Statement

All of the sequencing data generated in this study including the genome assembly generated for JF289 and small RNA sequencing data has been submitted to NCBI under the BioProject PRJNA996625. The genome assembly and annotations are also available via the figshare repository (DOI: 10.6084/m9.figshare.24975069).

