## Supplementary information for "Calcineurin contributes to RNAi-mediated transgene silencing and small interfering RNA production in the human fungal pathogen *Cryptococcus neoformans*"

**Supplementary Material and Methods**

***Scripts used for small RNA analysis***

Trim adapters using cutadapt:

~/.local/bin/cutadapt -j 8 --discard-untrimmed -m 14 -M 40 -a AACTGTAGGCACCATCAAT -g GTTCAGAGTTCTACAGTCCGACGATC sample.fastq -o sample-trimmed.fastq.gz

Map reads using bowtie2:

bowtie2 -x ref sample-trimmed.fastq.gz -S sample-trimmed.sam -p 8

Use Tim Dahlmann perl files for anlaysis and count reads from sam file along with nt preference analysis in a single step (<https://github.com/timdahlmann/smallRNA>)

perl /home/path/to/folder/count-read-length-from-sam.pl sample-trimmed.sam sample-reads.txt

Use the sample-reads.txt file for generating graphs in Excel or GraphPad Prism

***DEseq2 script used for differential analysis used***

\# Read the Geneious Output
###############################################################################
assayCsvFile - read.csv("assay.csv", row.names=1)
colDataFile - read.csv("colData.csv", row.names=1)
rowRangesFile - read.csv("rowRanges.csv")

###############################################################################
\# Prep for DESeq2
###############################################################################
assayMatrix - as.matrix( assayCsvFile )
\# having to do a transpose to get into the form we need: uses ludicrous memory to save in this format in Geneious
assayMatrix - t( assayMatrix )
colDataFrame - DataFrame( colDataFile, row.names=rownames( colDataFile ) )
rowRanges - makeGRangesFromDataFrame( rowRangesFile, start.field="start", end.field="end", starts.in.df.are.0based=FALSE )

countdata - assayMatrix
coldata - colDataFrame

\# Variables for use in design argument are columns found in colData
ddsFull - DESeqDataSetFromMatrix(countData = countdata, colData = coldata, design = ~ condition) # from count tables

\# gm_mean is from https://github.com/joey711/phyloseq/issues/387
gm_mean = function(x, na.rm=TRUE) {
exp(sum(log(x[x > 0]), na.rm=na.rm) / length(x))
}

\# taking rows not columns here (the 1 argument in apply function) because we transposed the counts earlier
eachGeneHasZeroes - all(!apply(countdata, 1, function(row) all(row != 0 )))

if(eachGeneHasZeroes) {
geoMeans - apply(counts(ddsFull), 1, gm_mean)
ddsFull - estimateSizeFactors(ddsFull, geoMeans = geoMeans)
}

###############################################################################
\# Post Processing
###############################################################################
dds - DESeq( ddsFull, fitType = "parametric" )
res - results( dds )

###############################################################################
\# Output Results
###############################################################################
outfilePath - paste(getwd(), "results.txt", sep="/")
write.table(as.data.frame(res), file = outfilePath, sep = "\t")

**Supplementary figures and figure legends**

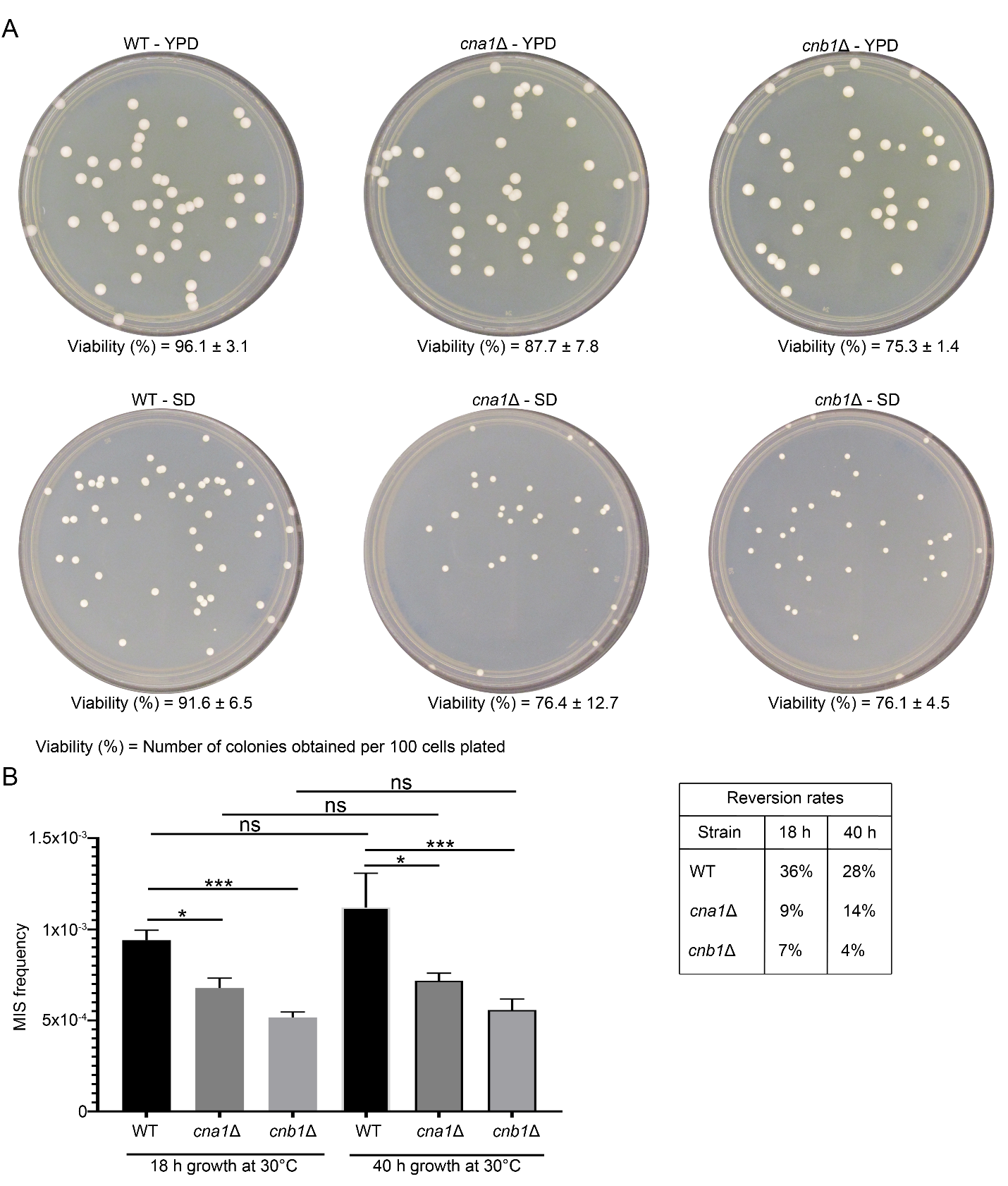

**Figure S1. Growth rate in calcineurin mutants does not impact their MIS frequency. (A)** Plate pictures showing the growth for the wild-type (JF289) and calcineurin mutants in both the nutrient-rich YPD media and nutrient limiting standard defined (SD) media. The viability percent is listed for each strain in both media conditions. **(B)** A graph showing the MIS frequency and a table presenting the MIS reversion rate in the wild-type (WT), *cna1*Δ and *cnb1*Δ mutants at two different time points from the same culture (p-values: * < 0.05, *** < 0.001, ns > 0.05).

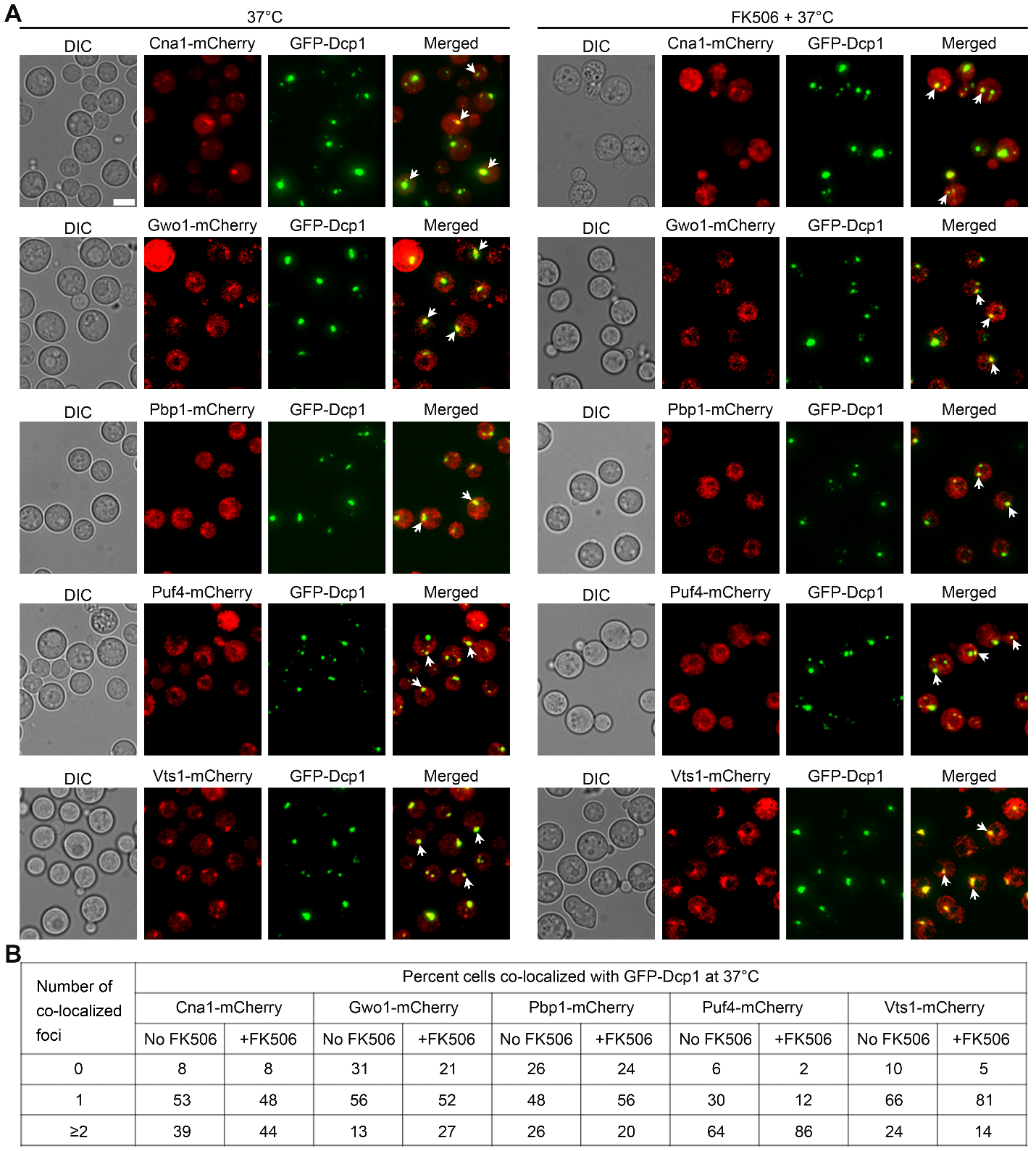

**Figure S2. Calcineurin activity is not required for the P-body localization of its substrates.** **(A)** Microscopy images showing the co-localization of the mCherry-tagged version of calcineurin and its substrates with GFP-Dcp1, a P-body marker, at 37°C in the absence and presence of calcineurin inhibitor, FK506. Bar, 5 µm. Some of the co-localization events are highlighted with white arrowheads in all cases. **(B)** A table describing the analysis of the number of cells showing co-localization with the P-body marker, GFP-Dcp1, in the presence and absence of FK506.

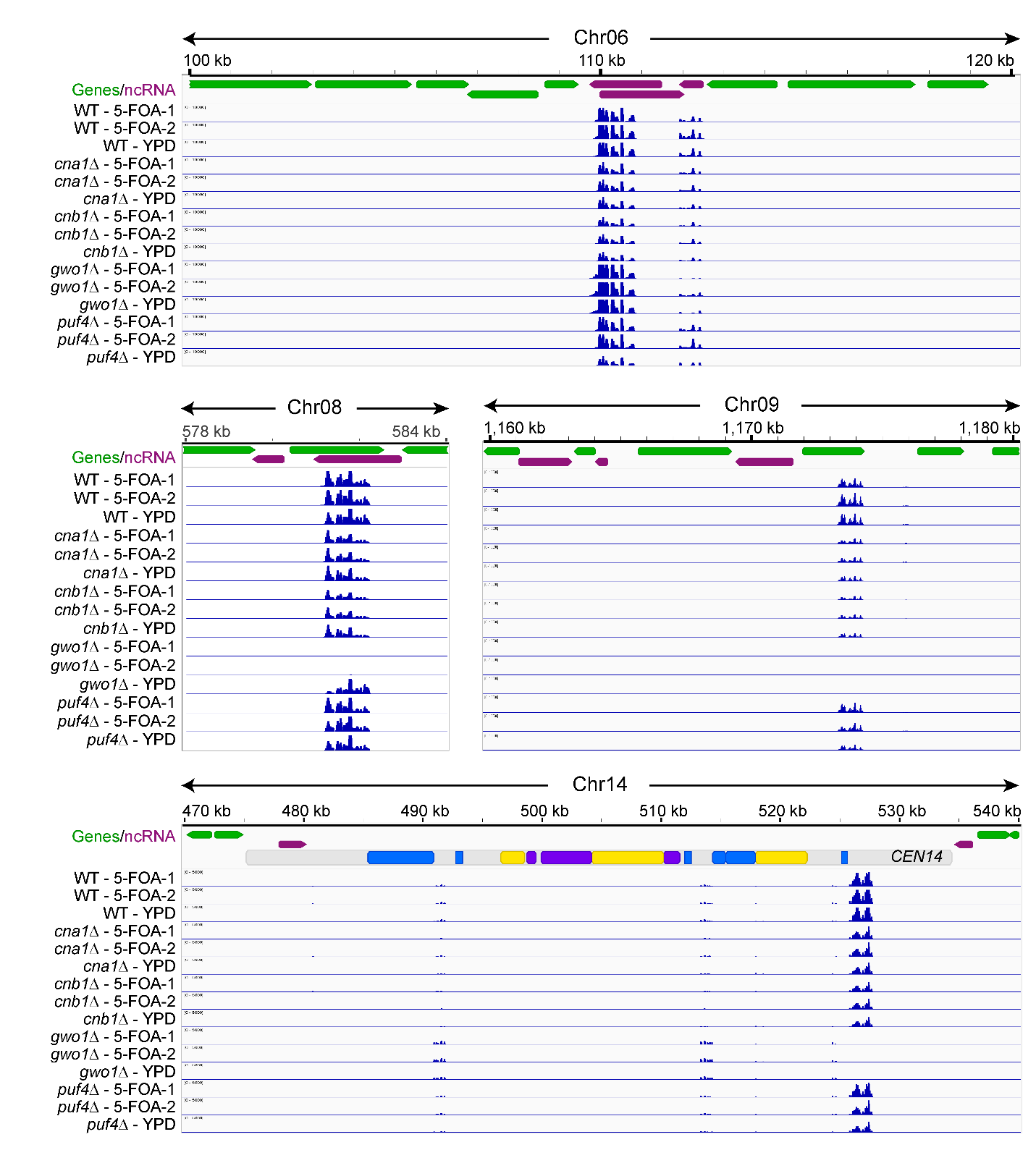

**Figure S3. Loss of calcineurin leads to reduced siRNA production across multiple genomic loci.** Small RNA reads mapping revealed a lower level of siRNA against multiple different regions of the genome in calcineurin mutants and a complete abolishment of siRNA against some genomic regions in the *gwo1*Δ mutant.

**Supplementary tables**

**Table S1. List of strains used in this study.**

| Strain name | Description | Reference |
| --- | --- | --- |
| JF289**a** | Wild-type *MAT***a** | (Wang *et al.* 2010) |
| H99 | Wild-type *MAT*α | (Perfect *et al.* 1993) |
| VYD225 | JF289**a** *cna1*Δ(*CNAG_04796*)::*NEO* | This study |
| VYD226 | JF289**a** *cnb1*Δ(*CNAG_00888*)::*NEO* | This study |
| VYD249 | JF289**a** *gwo1*Δ(*CNAG_00520*)::*NEO* | This study |
| VYD251 | JF289**a** *pbp1*Δ(*CNAG_02046*)::*NEO* | This study |
| VYD253 | JF289**a** *puf4*Δ(*CNAG_02810*)::*NEO* | This study |
| VYD255 | JF289**a** *vts1*Δ(*CNAG_06103*)::*NEO* | This study |
| VYD257 | JF289**a** *cna1*Δ::*HYG gwo1*Δ::*NEO* | This study |
| VYD259 | JF289**a** *cna1*Δ::*HYG puf4*Δ::*NEO* | This study |
| VYD261 | JF289**a** *puf4*Δ::NEO *gwo1*Δ::*NAT* | This study |
| HP1 | H99α *gwo1*Δ::*NEO* | (Park *et al.* 2016) |
| HP24 | H99α *vts1*Δ::*NEO* | (Park *et al.* 2016) |
| LK289 | H99α *CNA1*-*mCherry*::*NEO* *GFP-DCP1*::*NAT* | (Kozubowski *et al.* 2011) |
| WX250 | H99α GWO*1*-*mCherry*::*NEO* *GFP-DCP1*::*NAT* | (Park *et al.* 2016) |
| HP114 | H99α *PBP1*-*mCherry*::*NEO* *GFP-DCP1*::*NAT* | (Park *et al.* 2016) |
| HP130 | H99α *PUF4*-*mCherry*::*NEO* *GFP-DCP1*::*NAT* | (Park *et al.* 2016) |
| HP138 | H99α VTS*1*-*mCherry*::*NEO* *GFP-DCP1*::*NAT* | (Park *et al.* 2016) |

**Table S2. List of primers used in this study.**

| Primer name | Sequence (5’ – 3’) | Purpose |
| --- | --- | --- |
| JOHE50272 | CAGTAGGATCAAACACAATGGAAG | *CNA1* deletion construct |
| JOHE50273 | GCAAGGGCGAATTCTGCAGGACGGAAATTGACTGTTTGGTG |  |
| JOHE50274 | CACCAAACAGTCAATTTCCGTCCTGCAGAATTCGCCCTTGC |  |
| JOHE50017 | CGTTCGAAACCAGCATCTACTCCCAAGCTTGGTACCGAGCTC |  |
| JOHE50018 | GAGCTCGGTACCAAGCTTGGGAGTAGATGCTGGTTTCGAACG |  |
| JOHE50019 | GACATATCTAGCTCGTCTCACTC |  |
| JOHE50275 | CAAGAGTTAGCATGACTCACTTCG |  |
| JOHE50021 | CGCCGAGTCTGGATGGACAG |  |
| JOHE50276 | TGCCCTTCATTTGGATCATTG | *CNB1* deletion construct |
| JOHE50277 | GCAAGGGCGAATTCTGCAGTGCAATAAGGCGGTATTGATGATG |  |
| JOHE50278 | CATCATCAATACCGCCTTATTGCACTGCAGAATTCGCCCTTGC |  |
| JOHE50025 | CGTATATGGGGTAGGAATGAGAAACCAAGCTTGGTACCGAGCTC |  |
| JOHE50026 | GAGCTCGGTACCAAGCTTGGTTTCTCATTCCTACCCCATATACG |  |
| JOHE50027 | TGACGCCTCCTCCCAAGTC |  |
| JOHE50279 | CACCGTCGTACATTAGTATAATGG |  |
| JOHE50029 | TGGCTTTATTCTTCTCGCAGAG |  |
| JOHE51154 | ACCGGCAGGGTATACTGTTGCCGGGATAACGAGATTCGAGGTTTTAGAGCTAGAAATAGC | *CNA1* deletion guide RNA |
| JOHE51155 | ACCGGCAGGGTATACTGTTGGCAGTTGGAGAGGCTCGCAGGTTTTAGAGCTAGAAATAGC |  |
| JOHE51156 | ACCGGCAGGGTATACTGTTGAAGATGGGCGAGTCTCATGTGTTTTAGAGCTAGAAATAGC | *CNB1* deletion guide RNA |
| JOHE51157 | ACCGGCAGGGTATACTGTTGGAGATGTAAGTGTCAAGGCGGTTTTAGAGCTAGAAATAGC |  |
| JOHE52033 | TGAGTGGTGTAGTCAACGAGTG | *PBP1* ORF |
| JOHE52034 | GTTGATGTGGGGCGATAGAG |  |
| JOHE52035 | ACCGGCAGGGTATACTGTTGCCTCAGCATATGATGCCTGGGTTTTAGAGCTAGAAATAGC | *PBP1* deletion guide RNA |
| JOHE52036 | ACCGGCAGGGTATACTGTTGTACGACCAACAACATCACATGTTTTAGAGCTAGAAATAGC |  |
| JOHE52037 | CTCGTTCTCTCCCGTTAACG | *PUF4* ORF |
| JOHE52038 | CACATTCATGCTCAGAGCAAGG |  |
| JOHE52039 | ACCGGCAGGGTATACTGTTGCATGGTATGAGAGACCAGTGGTTTTAGAGCTAGAAATAGC | *PUF4* deletion guide RNA |
| JOHE52040 | ACCGGCAGGGTATACTGTTGTTAAGACCGATGGATCCAGGGTTTTAGAGCTAGAAATAGC |  |
| JOHE52041 | TCCAAGCTGTCCAATATGAACC | *VTS1* ORF |
| JOHE52042 | TCGATACTTGCGAGCATCGTC |  |
| JOHE52043 | ACCGGCAGGGTATACTGTTGGGAACCCTTTGTATAATCAGGTTTTAGAGCTAGAAATAGC | *VTS1* deletion guide RNA |
| JOHE52044 | ACCGGCAGGGTATACTGTTGAAAGTCCAACTGGAATGACAGTTTTAGAGCTAGAAATAGC |  |

**Table S3. Differential expression analysis of small RNA reads.**

Attached as a separate excel file.

**Table S4. Growth rates of wild-type and mutant strains in different media.**

| Strain | Doubling time in YPD  (mins ± standard deviation) | Doubling time in defined media  (mins ± standard deviation) |
| --- | --- | --- |
| JF289 | 109.8 ± 1.7 | 150.3 ± 6.6 |
| JF289 cna1 | 168.3 ± 11.6 | 195.6 ± 21.6 |
| JF289 cnb1 | 143.9 ± 2.6 | 212.8 ± 35.7 |
| JF289 gwo1 | 125.7 ± 2.6 | 162.0 ± 9.4 |
| JF289 pbp1 | 120.8 ± 9.4 | 156.2 ± 13.1 |
| JF289 puf4 | 140.1 ± 6.1 | 122.5 ± 20.6 |
| JF289 vts1 | 151.0 ± 6.4 | 110.3 ± 0.8 |
| JF289 rdp1 | 121.3 ± 7.6 | 107.7 ± 2.4 |
